# Sord deficient rats develop a motor-predominant peripheral neuropathy unveiling novel pathophysiological insights

**DOI:** 10.1101/2023.12.05.570001

**Authors:** Adriana P. Rebelo, Clemer Abad, Maike F. Dohrn, Jian J. Li, Ethan Tieu, Jessica Medina, Christopher Yanick, Jingyu Huang, Brendan Zotter, Juan I. Young, Mario Saporta, Steven S. Scherer, Katherina Walz, Stephan Zuchner

## Abstract

Biallelic *SORD* mutations cause one of the most frequent forms of recessive hereditary neuropathy, estimated to affect approximately 10,000 patients in North America and Europe alone. Pathogenic *SORD* loss-of-function changes in the encoded enzyme sorbitol dehydrogenase result in abnormally high sorbitol levels in cells and serum. How sorbitol accumulation leads to peripheral neuropathy remains to be elucidated. A reproducible animal model for SORD neuropathy is essential to illuminate the pathogenesis of SORD deficiency and for preclinical studies of potential therapies. Therefore, we have generated a *Sord* knockout (KO), *Sord*^−/−^, Sprague Dawley rat, to model the human disease and to investigate the pathophysiology underlying SORD deficiency. We have characterized the phenotype in these rats with a battery of behavioral tests as well as biochemical, physiological, and comprehensive histological examinations. *Sord*^−/−^ rats had remarkably increased levels of sorbitol in serum, cerebral spinal fluid (CSF), and peripheral nerve. Moreover, serum from *Sord*^−/−^ rats contained significantly increased levels of neurofilament light chain, NfL, an established biomarker for axonal degeneration. Motor performance significantly declined in *Sord*^−/−^ animals starting at ∼7 months of age. Gait analysis evaluated with video motion tracking confirmed abnormal gait patterns in the hindlimbs. Motor nerve conduction velocities of the tibial nerves were slowed. Light and electron microscopy of the peripheral nervous system revealed degenerating myelinated axons, de– and remyelinated axons, and a likely pathognomonic finding – enlarged “ballooned” myelin sheaths. These findings mainly affected myelinated motor axons; myelinated sensory axons were largely spared. In summary, *Sord*^−/−^ rats develop a motor-predominant neuropathy that closely resembles the human phenotype. Our studies revealed novel significant aspects of SORD deficiency, and this model will lead to an improved understanding of the pathophysiology and the therapeutic options for SORD neuropathy.

## Introduction

Biallelic mutations in *SORD* cause SORD deficiency, which is a common, recessive neuropathy that accounts for about 7-9% of distal hereditary neuropathy (dHMN) and Charcot-Marie-Tooth type 2 (CMT2) ^1^. The most common pathogenic variant, c.757delG (p.Ala253GlnfsTer27), has a population carrier frequency of 1/200-300 individuals in all major population ancestries. This variant alone is thus present in a homozygous state in approximately 1 per 100,000 individuals (∼3,000 in the US and ∼80,000 worldwide)^1^. *SORD* encodes the enzyme sorbitol dehydrogenase, a key enzyme in the polyol pathway, a two-step metabolic process that converts glucose to fructose ^2,3^. The first step of the pathway involves the conversion of glucose to sorbitol via aldose reductase, accompanied by the oxidation of NADPH to NADP+ ^4^. In the second step, sorbitol dehydrogenase catalyzes the metabolism of the sugar alcohol sorbitol to fructose, reducing NAD^+^ to NADH ^4^. Activation of the polyol pathway is triggered by excessive intracellular glucose, which is normally metabolized by hexokinases during glycolysis ^3^. During hyperglycemia, approximately 30% of glucose in the body passes through the polyol pathway, as an alternate route for glucose metabolism ^5^. Excessive production of sorbitol during hyperglycemia has been postulated to be an important mechanism of diabetic neuropathy ^6,7^. Moreover, increased activation of aldose reductase during hyperglycemia can lead to oxidative stress due to redox imbalance between NADH and NAD^+^ ^3,8^. Sorbitol accumulates in serum from patients with SORD deficiency due to loss of sorbitol dehydrogenase activity ^1^. However, distinct from diabetic neuropathy, the polyol pathway is not overactivated in SORD patients.

How sorbitol accumulation leads to neuropathy remains to be elucidated. High levels of sorbitol have been associated with several molecular mechanisms, including osmotic stress, toxicity, reactive oxygen species (ROS) production, and mitochondrial dysfunction ^9^. It is speculated that sorbitol does not readily diffuse across cell membranes and excess sorbitol causes increases intracellular osmolarity, leading to depletion of intracellular osmolytes ^10^. Like other neurons, motor neurons are non-insulin dependent cells, taking up a considerable amount of glucose in the absence of insulin and rendering them particularly vulnerable to the polyol pathway. In order to overcome sorbitol accumulation, aldose reductase inhibitors (ARIs) have been considered as a potential therapeutic approach for high sorbitol conditions. Consequently, a novel ARI is currently being tested in a clinical trial of SORD neuropathy (NCT05397665, clinicaltrial.gov). This oral drug crosses the blood brain barrier and showed significant reduction of serum sorbitol in SORD-deficient patients, in patient-derived fibroblasts and iPSCs-derived motoneurons, as well as in a *Drosophila Sord* model ^1,9,11,12^.

A reproducible animal model for SORD neuropathy is essential to study the mechanisms underlying the pathological abnormalities and to perform preclinical studies for potential drugs and gene therapy. A sorbitol dehydrogenase deficient mouse harboring a naturally occurring splice variant was reported to have no motor phenotype and no significant changes in motor nerve conduction velocity, despite developing abnormal accumulation of sorbitol ^13^. Mouse models failing to mimic human disease is a recurring issue, in particular in neuromuscular phenotypes. The laboratory rat has demonstrated to be a relevant and reliable model to study neurological human diseases, including CMT ^14,15^. They share physiological and genetic similarities with humans, making them valuable models for studying human diseases. Therefore, we generated a *Sord* knockout rat (Sprague Dawley), *Sord*^−/−^, to investigate in greater detail the neurological consequences of SORD deficiency. The *Sord*^−/−^ animal described herein exhibited abnormal accumulation of sorbitol and developed a behavioral motor-predominant neuropathy phenotype. This was demonstrated by electrophysiology, increased levels of NfL in serum, and histological studies of the PNS that showed changes predominately affecting large, myelinated motor axons. We also show previously unknown characteristics, such as even higher sorbitol accumulation in cerebral spinal fluid and ballooned myelin sheaths of motor axons. This *Sord^−/−^* model will be a valuable tool for mechanistic and therapeutic studies.

## Materials and methods

### Generation of the *Sord^−/−^* rat model

*Sord* knockout rats (in a Sprague Dawley genetic background) were generated at ENVIGO Inc. using CRISPR-CAS9 technology with a two guide RNA approach, each targeting the upstream and downstream coding regions of *Sord*, as indicated above. F1 animals were rats generated and genotyped at ENVIGO by PCR prior to shipment to the University of Miami. Briefly, genomic DNA was collected from a tail biopsy by adding 500 µl of TEL buffer and 10 µl of proteinase K 10mg/ml and incubated at 55°C ON with agitation. DNA fragments were amplified using primers Sord-F: CCCCTCTCTCTCCCTACCAG and Sord-R: GTTCTGCCTTTGGTCCACAT (for the wild-type allele) and US-F: GTGGTGGTGTGTTGAGCAAT and DS-R: ACTGCAGCCCAAGTTACACC (for mutant allele) and a protocol of 95°C for 2 min, then 35 cycles of 95°C for 45 s, 56°C for 1 min, and 72°C for 2 min, then a final 72°C for 5 min followed by 4°C until PCR products were visualized on an agarose gel. The expected band for the wild-type is 184bp, and the mutant is 399bp. Rats at the age of 2 months were shipped to the University of Miami and housed at a Specific Pathogen Free (SPF) facility with a 12 h light-dark cycle with access to food and water ad-libitum for the duration of the testing. All testing were performed in accordance with the National Institutes of Health Guide for the Care and Use and approved by the Institutional Animal Care and Use Committee at the University of Miami. In order to minimize number of animals and maximize the data obtained a comprehensive phenotyping scheme, described below, was followed. For electrophysiology and histological studies, animals from each genotype and sex were randomly selected.

### Behavioral studies

Behavioral testing began at 12 weeks of age and every two months thereafter until the animals reached 13 months of age. All testing was performed between 9 AM–1 PM. The initial number was 40 *Sord* ^−/−^ and 40 wild type (WT) rats, both males and females. The total number of individuals in each test is reported in the figure legends, and owing to the strategy and casual losses, the number of animals decreased over time. The testing battery lasted seven days and was performed blind to the genotype. A single test was performed daily in the following order: day 1: general characterization, days 2-4: narrow beam testing, days 4-7 Rota rod test, 8. hot plate test. Prior to each test, a habituation period of 30 minutes was provided in the testing environment. At the end of the tests animals were placed back in their home cage.

#### General characterization

The general health condition of the rats was evaluated by assessing their body weight, coat condition, and whiskers. The blink reflex response was measured followed to whiskers stimulation with a cotton swab.

#### Narrow beam

The narrow beam test was used to assess coordination. A 100 cm wooden beam (4 cm wide and 3 cm deep) was supported by tripods at both ends and elevated 80 cm off the ground. One end of the beam had a platform with a black box. Rats were placed at the other end, and the time to reach the opposite platform was recorded. Animals were subjected to 3 trials for two days for training and an extra day for testing.

#### Rota-rod test

A rotating rod was used to measure motor function and coordination in rats. The animals were placed on the accelerating rotarod apparatus (Ugo Basile) for nine trials over three consecutive days (three trials per day), with 15 minutes between each trial. The test consisted of two acceleration modes: the first mode started at 10 RPM and accelerated linearly to 40 RPM over a period of 100 seconds, and the second mode increased speed from 40 to 50 RPM for 200 seconds. The total trial time was 300 seconds. After two days of training, testing trials were conducted on the third day, and the time each rat took to fall from the rod, were recorded.

#### Hot plate test

The hot plate was used to detect any difference in pain perception. The response of a heat stimulus applied through the hind paw was recorded. Animals were placed in the center of the hot plate surface (Med Associate) heated to a maximum temperature of 55°C. The latency was recorded to indicate a nociceptive response with hind paw lick, flick, vocalization, or jump. The rat was removed once one of this reaction was observed. If there is no response within 30 seconds, the test is terminated.

### Gait Analysis

To assess the gait pattern of the rat model, we employed the MotoRater system (TSE Systems). The system consists of a narrow, transparent acrylic channel designed for mouse locomotion, with mirrors on both sides to capture different angles of motion. The system is integrated with a high-speed camera to track and record the rat’s movements. Various gait parameters were analyzed including the ankle angle extension, ankle angle flexion, ankle range of motion (ROM), and hip drop (maximum hip heigh minus minimum hip height during stride). To precisely locate these key points, we removed hair and marked each joint with a marker, which served as the reference points for the joints. Gait analysis was performed at 6-, 8-, 10– and 15-month-old rats (n=3 for each age group with the exception of the 15 months old rat, n=1). Raw data was collected and analyzed with the MotoRater software. Statistical analysis was performed with Graph Pad Prism. An unpaired Multiple Mann-Whitney test was performed with a False Discovery Rate (FDR) approach using the two-stage linear step-up procedure of Benjamini, Krieger and Yekutieli.

### Nerve conduction studies

We used the Nicolet EDX portable device (Natus) to measure nerve conductions. Rats were anaesthetized with isoflurane and paw temperature was maintained between 29.5-33.2°C. For stimulation and measurement, we used superficial ring electrodes for the tail and needle electrodes for the sciatic and tibial nerve positioned at standard positions. The grounding electrode was a clip electrode held at the opposite paw. Nerve conduction velocities (m/s) were calculated subtracting latencies from the two stimulation sites at the sciatic notch and lateral to the ipsilateral Achilles tendon, divided by the individually measured distance.

### Sorbitol measurement

Sorbitol measurements from serum, cerebral spinal fluid (CSF), and tissues were performed by UPLC–tandem mass spectrometry (MS/MS) (Waters Acq UPLC–tandem mass spectrometry (MS/MS) (Waters Acquityity UPLC & TQD mass spectrometer). Blood was collected after overnight fasting in serum separator tubes. Samples were centrifuged at 500g for 10 min, and serum was separated and frozen within 1 h from blood collection. Rat tissues were lysed in RIPA buffer (ThermoFisher) in the absence of protease inhibitor, which contains mannitol, a sorbitol enantiomer that can interfere with UPLC–MS/MS sorbitol determination. Samples underwent protein precipitation with acetonitrile (1:5), tenfold dilution with acetonitrile/water (50:50) and cleanup on Oasis HLB cartridges (10 mg/1 ml), before injection (3 μl) into the UPLC system. The UPLC conditions were as follows: column, BEH amide 1.7 μm (2.1 × 100 mm) at 88°C; eluent A, acetonitrile 90%/water 5%/isopropanol 5%; eluent B, acetonitrile 80%/water 20%; gradient elution, 0 min 100% A to 3.6 min 100% B; flow rate, 0.45 ml/min. The retention time of sorbitol was 2.7 min. The linearity of the method was assessed between 0.25 and 50 mg/l. The MS/MS conditions were as follows: interface, electrospray interface in negative ion mode; multiple reaction monitoring acquisition, m/z 180.9 → 88.9 (CV 24, CE 15). The detection limit (signal-to-noise ratio = 3) was 0.03 mg/l. The sorbitol level was tested by UPLC using a method adapted from Cortese et al^1^. The conditions were as follows: column, BEH amide 1.7 μm maintained at 25 °C (instead of 45 °C); eluent A, 10 mM ammonium acetate pH 10; eluent B, acetonitrile; flow rate, 0.6 ml/min with the same gradient. The retention time of sorbitol was 6.0 min. The MS/MS conditions were the same as for the fibroblast analysis ^1^. The serum samples underwent protein precipitation with cold methanol (1:5), fivefold dilution with acetonitrile/water (50:50) and cleanup on Oasis HLB cartridges (10 mg/1 ml), before injection (3μl) into the UPLC system. The calibration curve was prepared in serum in the sorbitol concentration range 0.1–20 mg/l.

### Light and electron microscopy

*Sord*^−/−^ rats (n=6) and their WT littermates (n=6), 64-75 weeks of age, were euthanized with an intraperitoneal injection of ketamine (200 mg/kg) and xylazine (10 mg/kg), then transcardially perfused with 50 ml of 0.9% NaCl followed by 300 ml of 2% paraformaldehyde and 2% glutaraldehyde in 0.1 M phosphate buffer (PB; pH = 7.4). The sciatic nerves (at the sciatic notch and at the knee), the tibial nerve at the ankle, the femoral motor and sensory nerves, and the 5^th^ lumbar root were dissected and fixed for 3 hours, then rinsed in 0.1M PB, transferred to a 2% OsO4 in 0.1M PB, for 1 hour, then processed for embedding in Epon ^16^. Semi-thin sections were stained with alkaline toluidine blue and visualized by light microscopy (Leica DMR) using interactive software (Leica Application Suite). For electron microscopy, transverse sections were cut at a thickness of 90 nm, and stained with lead citrate and uranyl acetate, and imaged using a JEOL 1010 electron microscope. Selected images were processed with Photoshop to generate the figures.

### Quantification of anatomical findings

Pathological features were quantified from semi-thin sections of the femoral motor nerve as previously described {Abrams, 2023 #34072}. To measure axonal calibers and g-ratios, EM images (2000X) from non-overlapping fields were taken from transverse sections of femoral motor nerves (3 *Sord* KO and 3 WT). All intact myelinated fibers, excluding regenerative clusters, were traced using Adobe Photoshop. g-ratios were calculated by the square root of the ratio of inner to outer traced axonal area using a custom script in Microsoft Excel with a circularity threshold of > 0.3 and a diameter threshold of > 4 μm. Graphed Prism was used for statistical analysis and figure generation.

### Simoa NFL assay

NfL was measured in rat serum using the Quanterix Simoa® NfL Assay following the manufacturers recommendations.

## Statistics

GraphPad Prism 10 (GraphPad Software, San Diego, CA, USA) was used for statistical analyses and figure generation for sorbitol concentration, NfL, nerve conduction studies, g-ratio quantification, behavioral analysis and gait analysis. For gait analysis, an unpaired Multiple Mann-Whitney test was performed with a False Discovery Rate (FDR) approach using the two-stage linear step-up procedure of Benjamini, Krieger and Yekutieli.

Sorbitol and NfL concentrations, Nerve conduction studies, as well as behavioral tests data were analysed using either a two-way ANOVA, or an unpaired *t*-test. All ANOVA main effects were followed by Tukey *post hoc* tests.

## Data availability

The corresponding author has full access to data presented in this article and it is available upon request.

## Results

### Generation of *Sord* knockout rats

A *Sord* knockout rat (*Sord*^−/−^; KO) was generated by CRISPR/Cas9-mediated deletion of the *Sord* gene. Two sgRNAs were designed to target two sites, one in exon 2 and another one in the 3’-UTR, genomic coordinates (approximately at chr3:114,188,869 and chr13:114,207,088) to generate a ∼18,238 bp deletion between the cleaved sites (Fig. 1a). As shown in Fig. 1b-c, animal #16 carried the expected *Sord* deletion, which was confirmed by Sanger sequencing (Supplementary figure 1). The founder animal was backcrossed to generate F1 heterozygous progeny, which were intercrossed to produce homozygous *Sord*^−/−^ rats. A Western blot demonstrated that sorbitol dehydrogenase protein was completely absent in the brain, spinal cord, and sciatic nerves of *Sord*^−/−^ rats (Fig. 1d). The level of aldose reductase protein was higher in sciatic nerve compared to brain and spinal cord, suggesting a high flux demand of glucose through the polyol pathway in the peripheral nerve.

**Figure 1:**
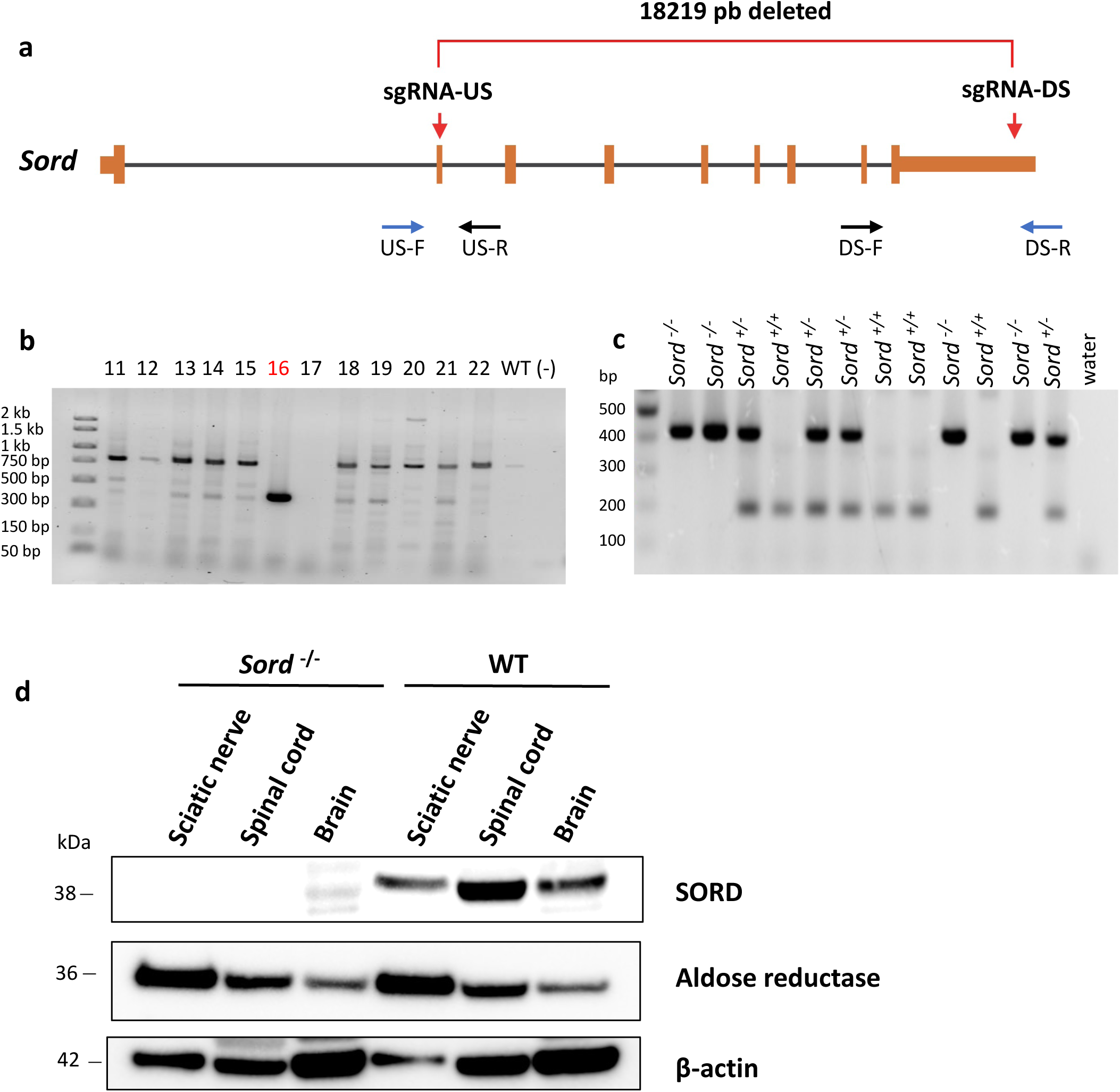
Generation of *Sord*^−/−^ rat and confirmation of loss of sorbitol dehydrogenase. (a) Diagram depicting the rat *Sord* gene and CRISPR-Cas9 gRNA target sites at exon 2 and the 3’-UTR. Primers flanking each sgRNA site (red arrows) were designed to test individual NHEJ activity, as well as paired together to screen for deletion mutations between the two target sites. (b) PCR amplification of genomic region spanning the gRNA target sites in rats treated with CRISPR-Cas9. Rat #16 contained a deleted region 18,238bp in the *Sord* gene. Expected mutation band using US-F and DS-R primers: ∼399bp (c) A representative result of genotyping PCR showing rats with heterozygous (e.g. lane 507) and homozygous (e.g. lane 505) deleted as well as unmodified (e.g. lane 508) *Sord.* (d) Western blot measuring Sord and aldose reductase protein in different tissues isolated from WT and *Sord*^−/−^ rats. Sord protein is absent in the *Sord*^−/−^ rats.

### Increased levels of sorbitol in the *Sord* deficient rats

Sorbitol levels are an important biomarker and diagnostic tool for SORD neuropathy^1^. We therefore measured sorbitol in rats by ultra-performance liquid chromatography-tandem mass spectrometry (UPLC/MS). Sorbitol was measured in serum, CSF, and in tissues isolated from 14 months old rats. Similar to the SORD deficient patients, sorbitol was significantly higher (∼7×) in *Sord*^−/−^ rats serum compared to WT siblings (Fig. 2a). Surprisingly, and previously unrecognized, sorbitol was 30× higher in the CSF of *Sord*^−/−^ rats compared to WT rats, indicating that neuronal tissues produce high levels of sorbitol via the polyol pathway (Fig. 2b). Additionally, sorbitol was highly elevated in homogenates of the spinal cord, sciatic nerve, and to a lesser extent in the kidney isolated from *Sord*^−/−^ rats (Fig. 2c). These findings confirm pronounced sorbitol accumulation in the peripheral nerve. Because glucose is the sole substrate for producing sorbitol via aldose reductase, we tested whether a high-glucose diet could further increase sorbitol levels. For this, 7-month-old *Sord*^−/−^ rats were placed on a high glucose (10%) diet for 2 months, but this did not significantly affect serum sorbitol levels compared to *Sord*^−/−^ rats on a regular diet (Supplementary figure 2), suggesting that the standard rat diet contains enough glucose to produce maximum sorbitol levels in serum.

**Figure 2:**
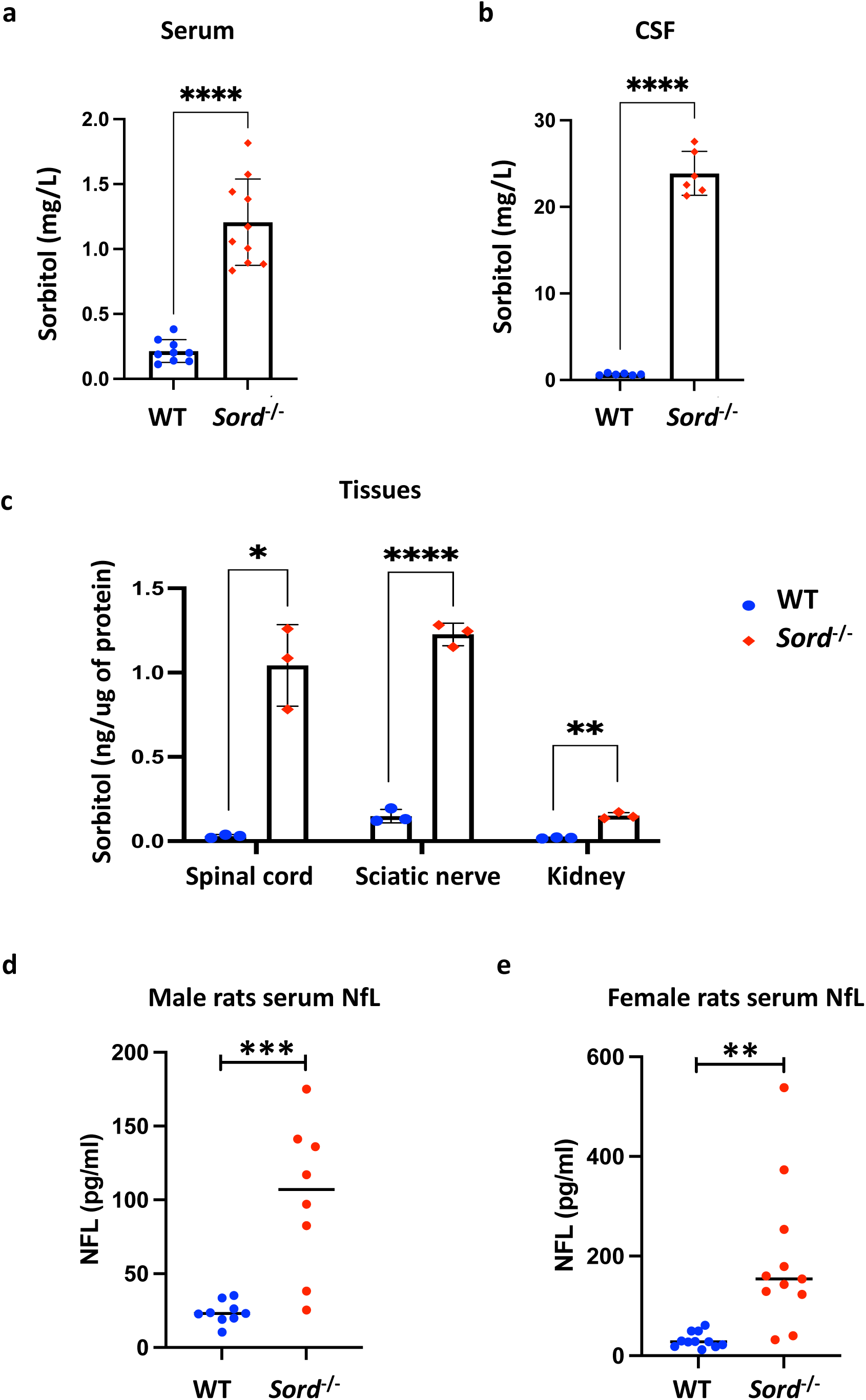
Biochemical studies performed in *Sord*^−/−^ rat samples. (a-c) Sorbitol levels measured by MS/MS in serum (a), CSF (N=10 each genotype)(b) and different tissues (N=6 each genotype)(c) isolated from 14-month-old rats. (N=3 each genotype) (d-e) NFL levels measured in serum of 15-month-old male (d) and female (N=11 each genotype) (e) rats by Simoa NF-light assay.

### Elevated NFL levels in serum from *Sord*^−/−^ rats

The serum/plasma level of neurofilament light (NfL), a cytoskeleton protein highly expressed in axons, has been used as a biomarker for axonal degeneration in neurological diseases, including CMT ^17^. We measured serum NfL in rat serum and found significantly increased levels of NfL in both male and female *Sord^−/−^* rats at 15 months of age compared to age-matched WT animals (Fig. 2d and Fig. 2e). These results suggest that axons are continuously damaged in *Sord*^−/−^ rats, and it raises the possibility that NfL could serve as a treatment-responsive biomarker.

### Motor deficits in *Sord*^−/−^ rats

*Sord*^−/−^ rats developed normally without any apparent phenotype. The body weight of the *Sord*^−/−^ animals, recorded at different ages, was comparable to the WT animals (Fig. 3a). We subjected a total of 10 rats to a battery of behavioral tests to measure motor coordination and muscle strength including rotarod, narrow beam test, grip strength, and gait. Young *Sord*^−/−^ animals (3-5 months of age) showed no significant difference in rotarod performance compared to WT rats (Supplemental Figure 3). However, motor performance of the adult male *Sord*^−/−^ rats started to progressively decline beginning at 7 months of age as revealed by their rotarod performance (Supplementary Figure 3). At 11 months of age, male *Sord*^−/−^ animals exhibited significantly shorter times to fall off the rotarod (latency time) compared to WT (Fig. 3b). The rotarod motor deficit was even more pronounced at the age of 13 months, suggesting a progressive motor deterioration with aging. Rats were also tested on a horizontal narrow beam for further fine coordination and balance evaluation. Both male and female *Sord*^−/−^ rats showed a decline in performance in the narrow beam test starting at 9 months of age. The average time for *Sord*^−/−^ rats to cross the narrow beam was significantly increased compared to WT (Fig 3c). Male *Sord*^−/−^ rats appear generally more affected than females in both rotarod and narrow beam tests. However, prior studies have demonstrated that female rats tend to significantly outperform male rats in the rotarod test ^18^

**Figure 3:**
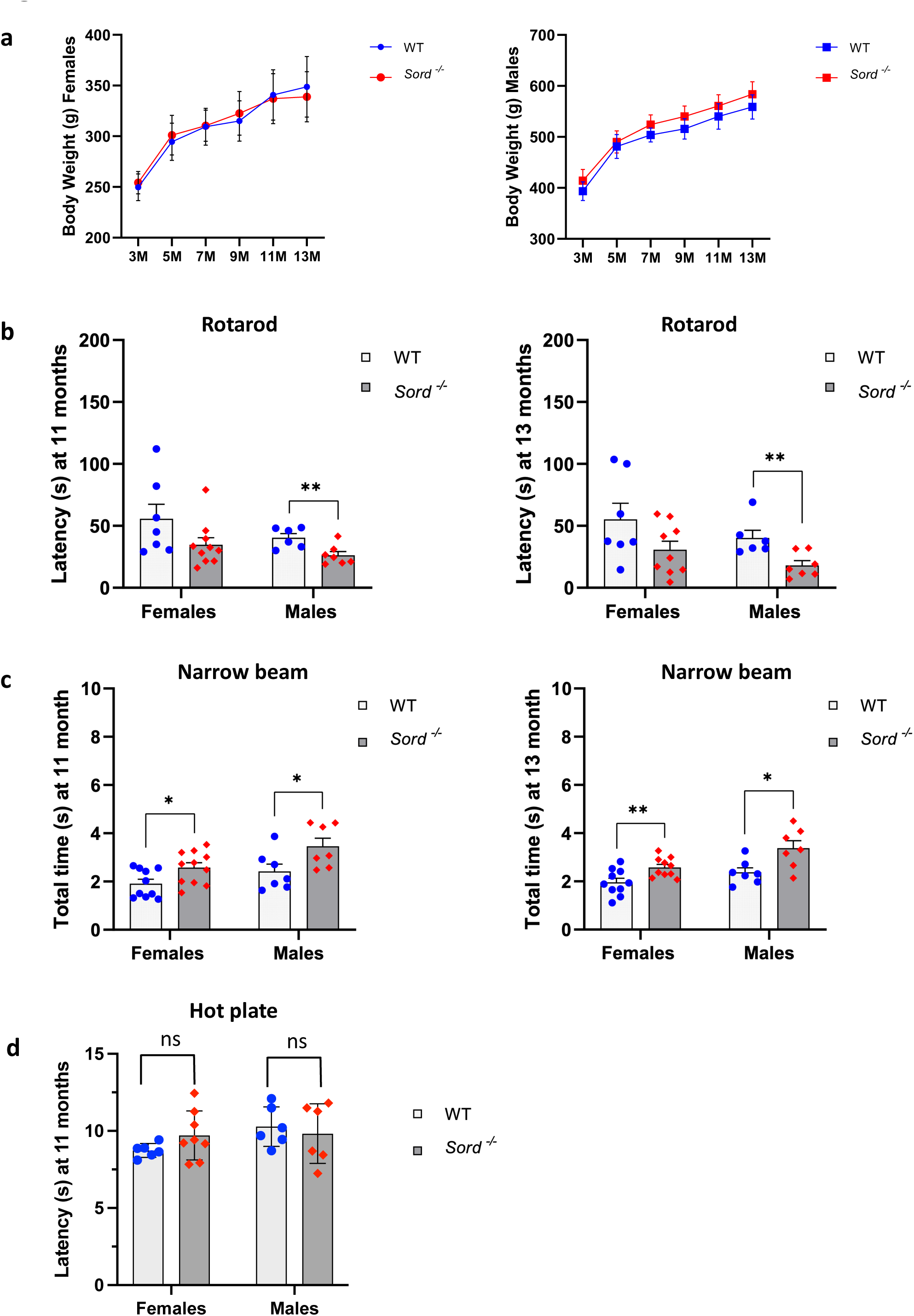
Assessment of motor and sensory function of *Sord*^−/−^ rats. (a) Body weights of male and female rats recorded at different ages (months). (b) Rotarod test show significantly decreased latency time of male *Sord*^−/−^ rats at both, 11 and 13 months of age. Latency time is recorded as the time it takes for the animals to fall off the rod. No significant differences were observed between females *Sord*^−/−^ and WT rats. (c) Narrow beam test shows significantly increased time for the *Sord*^−/−^ rats to cross from one end to the other end of the beam. (d) Hot plate test shows no differences in time to respond to noxious heat between WT and *Sord*^−/−^ rats. Data are presented as average +/− SEM. *= *P*<0.05, n=8 for each group.

### Normal sensitivity to heat in *Sord*^−/−^ rats

Diabetic neuropathy patients exhibit increased pain sensitivity in the extremities. Thus, we evaluated whether nociception is altered in the *Sord*^−/−^ rats. We performed the hot plate test to assay the latency to react to noxious heat at 11 months of age. *Sord*^−/−^ showed no significant difference in reaction times in the hot plate test, indicating normal nociception and suggesting that the motor phenotype is not accompanied by a significant sensory neuropathy at this age (Fig. 3d).

### Gait Analysis of *Sord*^−/−^ rats

For a more precise motor evaluation, gait analysis was performed on different age groups with the MotoRater, an automated system for fine motor kinematic video analysis. The system consists of a narrow channel with mirrors on both sides to monitor different angles of movement and a high-speed camera (Supplementary video). Five points were marked on the animal joints to track their movement during walking. Results revealed significant alterations in the ankle movement in the older *Sord*^−/−^rats, starting at 8 months of age. At 15 months of age, the gait abnormality in the *Sord*^−/−^ rats was more evident (Supplementary Video), especially more pronounced on the hindlimbs. The ankle flexion and ankle extension angles were significantly decreased in the older *Sord*^−/−^ rats compared to the WT (Fig. 4a and 4b), resulting in a significantly increased ankle range of motion in the *Sord*^−/−^ rats (Fig. 4c). These results are similar to previously reported gait patterns of a CMT1A mouse model ^19^. Moreover, the ‘hip drop’ height was significantly increased (Fig. 4d). These results confirmed impaired motor performance of *Sord^−/−^*rats.

**Figure 4:**
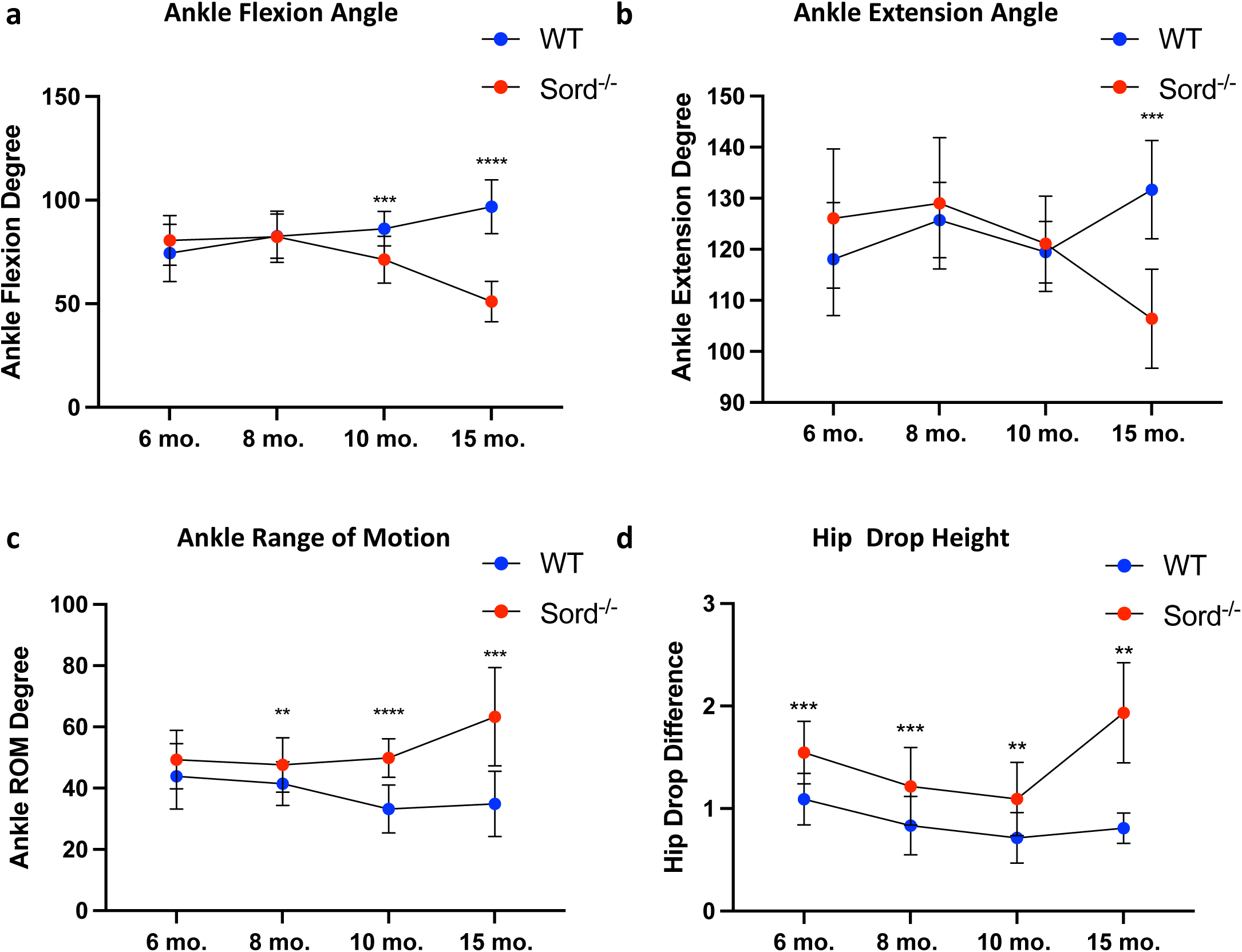
Gait analysis performed with the MotoRater instrument at 6-, 8-, 10– and 15-month-old rats. (a) Ankle flexion angle. (b) Ankle extension angle. (c) Ankle range of motion. (d) Hip drop height. Statistics: Unpaired Multiple Mann-Whitney test was performed with a False Discovery Rate (FDR) approach using the two-stage linear step-up procedure of Benjamini, Krieger and Yekutieli (n=3 for each age group with the exception of the 15 months old rat, n=1).

### Reduced nerve conduction velocity in *Sord*^−/−^ rats

In order to assess the peripheral nerve function, we performed nerve conduction studies on the sciatic, tibial, and caudal nerves. Using the same methodological concept as in humans, we measured compound muscle action potentials (CMAP) and calculated motor nerve conduction velocities (NCV) between tibial stimulation sites in 13 months old rats. A moderate, but significantly decreased sciatic NCV was observed in the *Sord^−/−^* male rats, but not in female rats (Fig. 5a-5d), indicating dysfunction of the motor axons. Sciatic and caudal nerve action potentials did not show significant differences compared to the WT. These findings suggest altered myelination and/or loss of large, myelinated motor axons in *Sord*^−/−^ rats.

**Figure 5:**
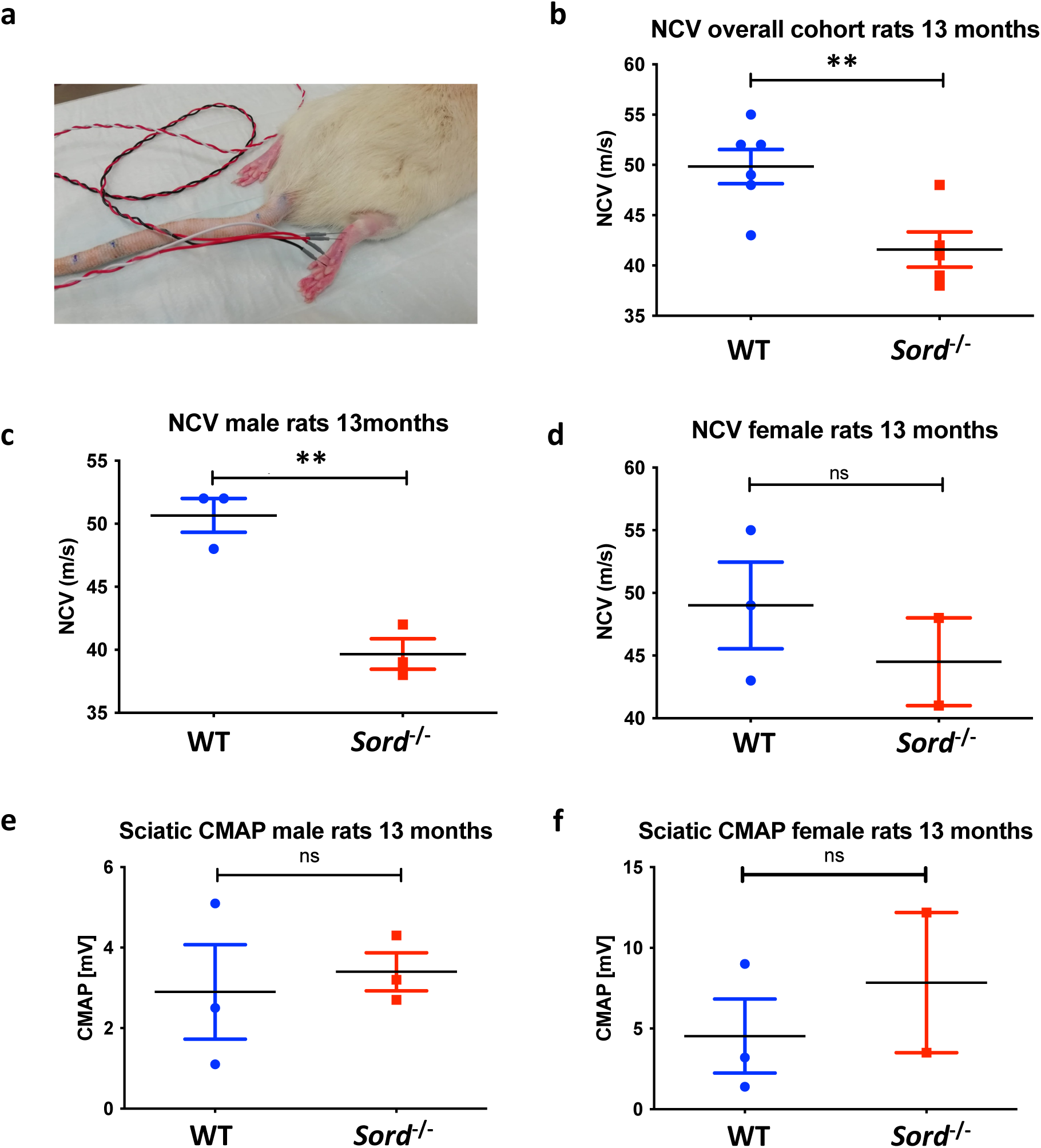
Electrophysiological studies performed in the peripheral nerve of *Sord*^−/−^ rats. (a) Image of Sord rat undergoing EMG studies. (b-d) Nerve conduction velocity (NCV) recorded from rat tibial nerve in 13-month-old rats – overall cohort (b), males (c), and females (d) Note that male *Sord* KO rats show significantly decreased NCV compared to WT, but no significance was observed in female rats (d). (e-f) Animals show no significant change of measured CMAP (n=3 for each sex group).

### Histological and ultrastructural abnormalities in the peripheral nerve of the *Sord*^−/−^ rats

We examined selected aspects of the peripheral nerves from 64–85-week-old *Sord*^−/−^ and WT rats – the ventral and dorsal roots, femoral sensory and motor nerves, the sciatic nerve at the hip and knee, the tibial nerve at the ankle, and the caudal nerve. Results are illustrated and quantified in Figures 6-8 and Supplementary Figures 4-5. The main findings were limited to large, myelinated axons in motor nerves (ventral roots, femoral motor, peroneal, and sciatic/tibial nerves) and observed less in sensory nerves (dorsal roots, femoral sensory, sural, and caudal nerves). A loss of large fibers and other features were present at all studied levels of the peripheral nervous system but appeared more pronounced in the distal sections (i.e., ventral root vs distal tibial nerve sections). The most severely affected areas of peripheral nerves showed mild endoneurial edema (Fig. 6 and Supplementary Fig. 4). The most common findings were degenerated axons, clusters of regenerating axons, thinly myelinated axons, and occasional de– and remyelinating axons with “onion bulb”-like formations with few layers of circumferential Schwann cell processes. An unusual yet consistent observation was abnormally enlarged (“ballooned”) myelin sheaths that still surrounding intact axons (Figs. 6 and 7c). Electron microscopy studies showed a “bubbly” disintegration of a compact myelin sheath revealing large structureless spaces between lamellae with the axon inside. Similar changes were described in rat galactose neuropathy models and a feline model of diabetic neuropathy and were associated with alterations in the polyol pathway ^20–22^.

**Figure 6:**
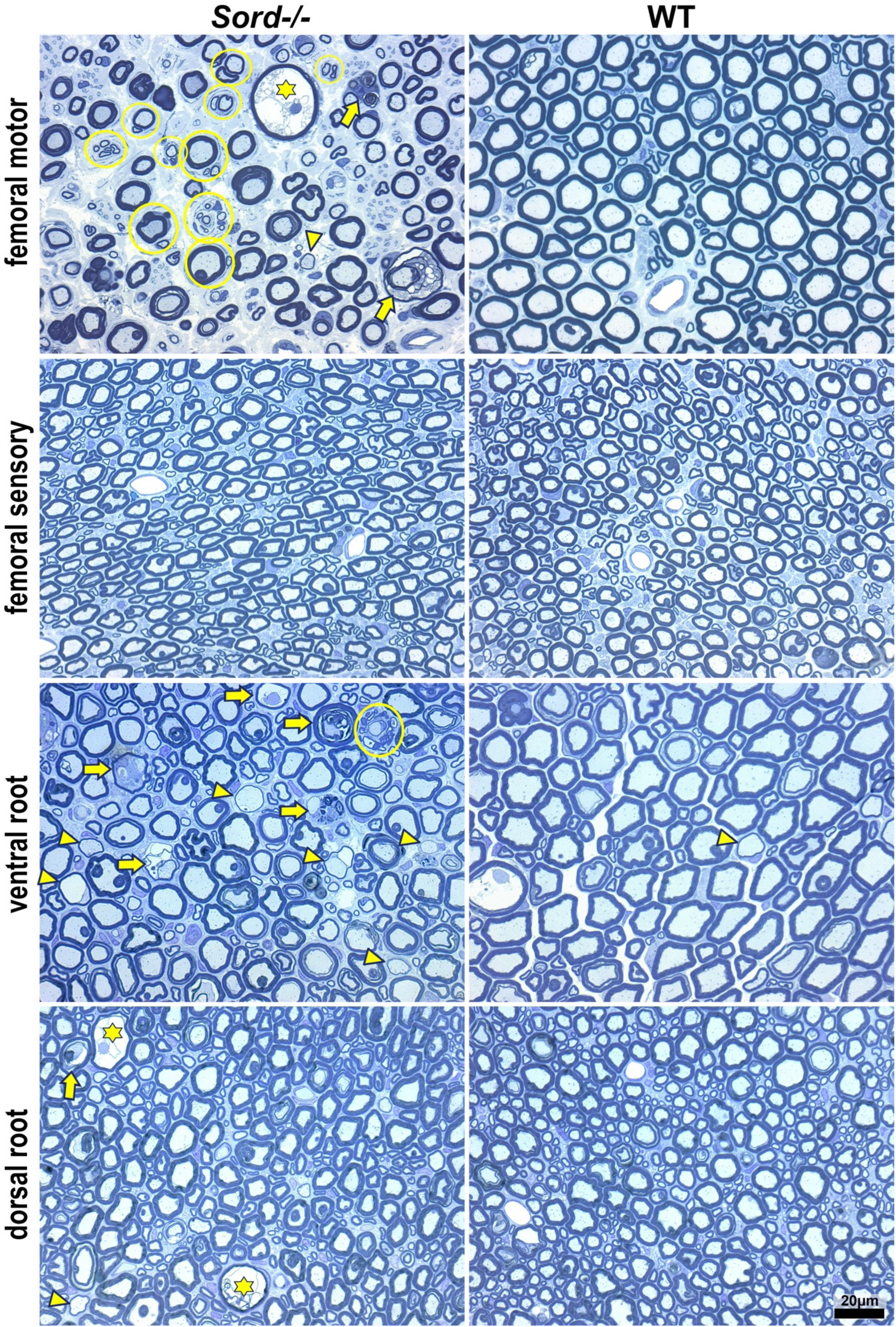
Myelinated motor axons are more affected than myelinated sensory axons in *Sord*^−/−^ rats. Representative images of semi-thin sections of the femoral motor, femoral sensory, ventral root, and dorsal root, from *Sord*^−/−^ and wild type (WT) rats, as indicated. The pathological findings are far more pronounced in nerves that contain myelinated motor axons, including degenerating myelinated sheaths (arrows), thinly myelinated axon (arrowheads), clusters of regenerated axons (circled), and “ballooned” myelin sheaths (star). The sensory branch of the femoral nerve is largely preserved; however, few “ballooned” fibers are observed in the dorsal root. Scale bar = 20 µm. *Sord*^−/−^ rats (n=6) and their WT littermates (n=6),

**Figure 7:**
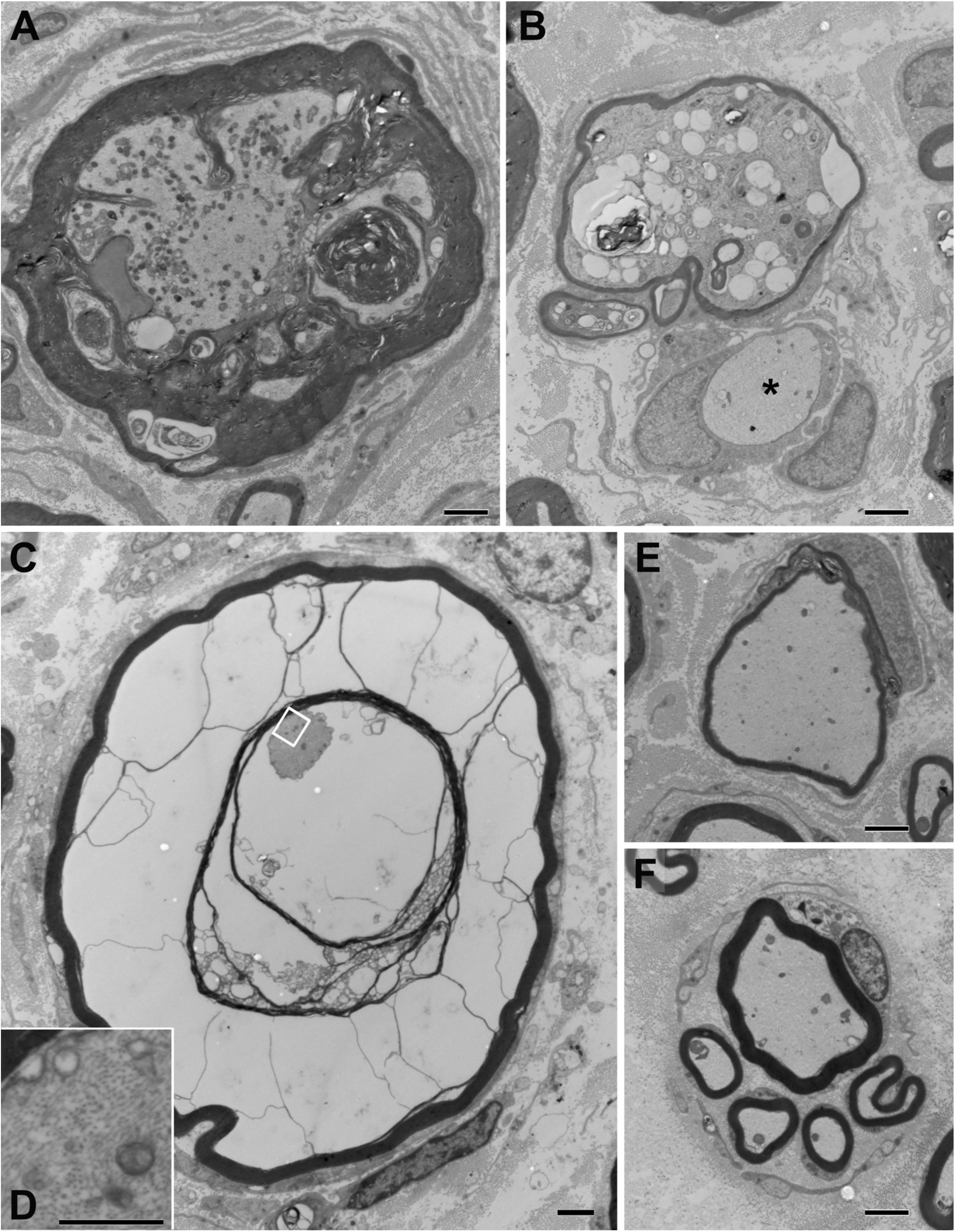
Ultrastructural features of myelinated axons in *Sord*^−/−^ rats. Electron micrographs of thin sections of femoral motor nerves from *Sord*-null rats illustrating degenerating myelinated sheaths (a and b), demyelinated axons (asterisk in b), thinly myelinated axons (e), clusters of regenerated axons (f), and “ballooned” myelin sheaths (c); inset (d) shows a magnified portion of the axon. Scale bars: 2 µm (a, b, c, e and f); 500 nm (d). Animals analyzed: *Sord*^−/−^ rats (n=6) and their WT littermates (n=6).

**Figure 8:**
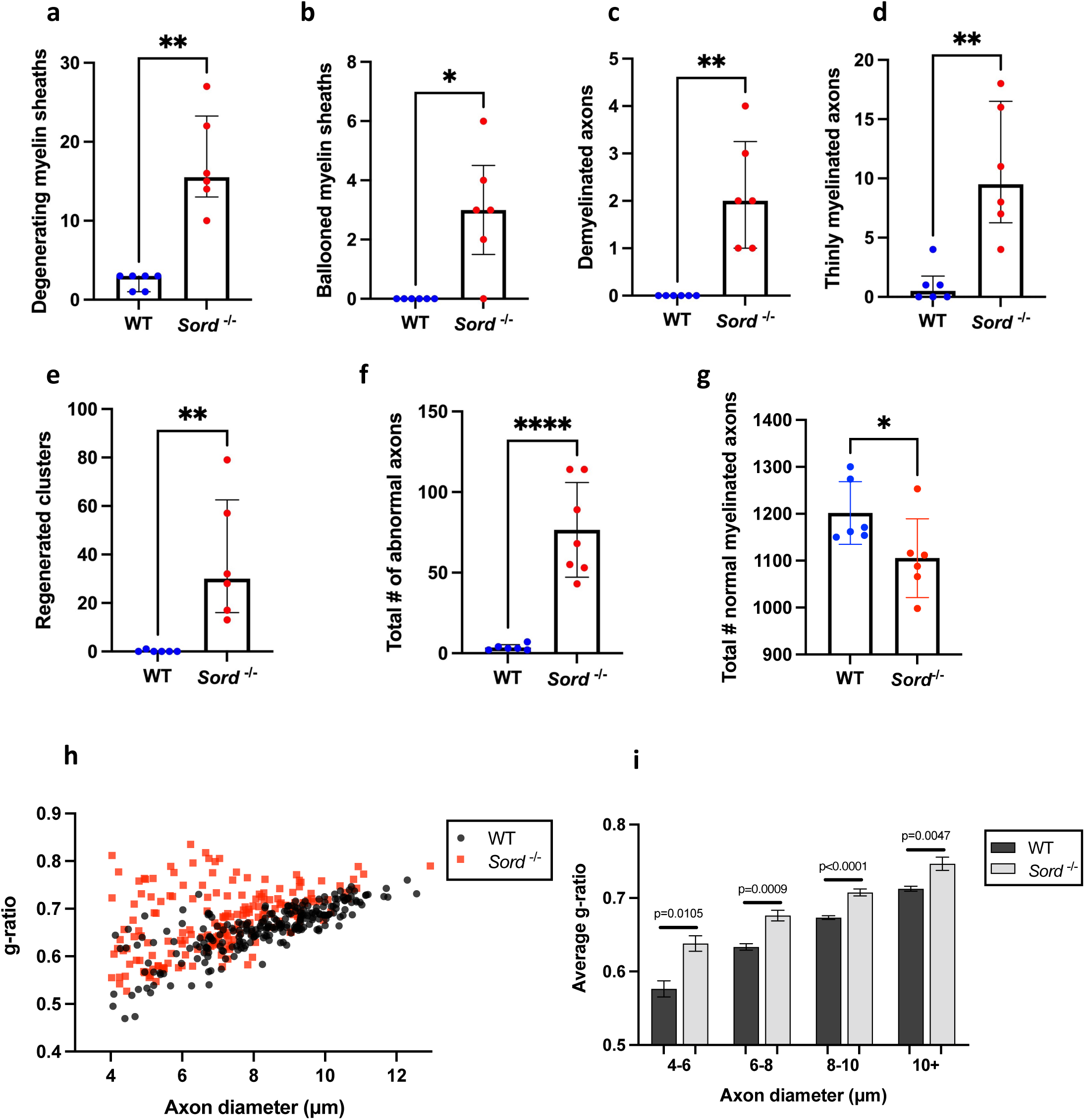
Quantification of pathological features observed in the *Sord*^−/−^ rats femoral motor nerves. The graphs show the significantly more degenerating myelin sheaths (a), ballooned myelin sheaths (b), demyelinated axons (c), thinly myelinated axons (d), clusters of regenerated axons (e), and the total number of axons with one of these abnormalities (f) in individual *Sord*^−/−^ rats (n=6) as compared to WT rats (n=6) (g) The total number of myelinated axons was significantly (Mann-Whitney U test). Each point represents a single animal. (h, i) Hypomyelination of medium and large axons in adult *Sord^−/−^* femoral motor nerves. (h) The g-ratios of myelinated axons > 4 μm was significantly increased in *Sord*^−/−^ (n=176 axons; from 3 rats) nerves, 70-85 weeks of age) as compared to 18-month-old WT (n=236 axons; from 3 rats) nerves (p<0.0001), and for axons binned by diameter and g-ratios were significantly increased across all bins in *Sord^−/−^* vs WT, p values as displayed. Error bars +/− SEM. The Kolmogorov–Smirnov test was used to determine significance.

We quantified axon and myelin pathology in the femoral motor nerves of 70-83 week-old *Sord*^−/−^ and WT rats. Numbers of degenerating myelin sheaths, “ballooned” myelin sheaths, demyelinated axons, thinly myelinated axons, and regenerated clusters, as well as the total number of abnormal axons, were significantly higher in *Sord*^−/−^ (n=6) compared to WT (n=6) rats (Fig. 8a-f). The total number of myelinated axons was reduced in *Sord*^−/−^ (n=6) compared to WT (n=6) rats (Fig. 8g; p=0.31, Mann-Whitney U test). We plotted the diameter and g-ratio of myelinated fibers from electron micrographs of *Sord*^−/−^ (n=3) and WT (n=3) rats (Fig. 8h). Axon diameters were significantly smaller (p = 0.0001; Kolmogorov–Smirnov test) in *Sord*^−/−^ (n=3) compared to WT (n=3) rats. G-ratios were significantly greater, both considering the population of myelinated axons as a whole (p = p<0.0001; Kolmogorov– Smirnov test), as well as when myelinated fibers were binned by diameter (Fig 8i) in *Sord*^−/−^ (n=3) compared to WT (n=3) rats. We did not determine whether these thinner myelin sheaths are a consequence of development, de/remyelination, or myelination of regenerated axons.

## Discussion

The anatomical and physiological complexities of the peripheral nerve pose challenges to *in vitro* studies. Thus, animal models of monogenic inherited peripheral neuropathies have long been valuable research tools ^23,24^. This increasingly includes the pre-clinical evaluation of gene-targeting, rational therapeutic interventions. SORD neuropathy was only recently described but is likely the most common recessive form of axonal inherited neuropathy, including the distal hereditary motor neuropathies. Moreover, sorbitol dehydrogenase is affected by a loss-of-function mechanism, rendering the outflow of the polyol pathway inactive. The lack of neuropathy in *Sord* deficient mice motivated us to develop a rat rodent model. In this study, we have generated and characterized a *Sord* gene knockout rat model, *Sord^−/−^*.

Our comprehensive analysis of *Sord*^−/−^ rats revealed a phenotype that closely resembles the findings in SORD neuropathy patients. This includes firstly the elevated levels of sorbitol in serum and urine, but also in different tissues such as the sciatic nerve. Intriguingly, we show that CSF from *Sord^−/−^* animals contains even higher levels of sorbitol compared to serum. This is in agreement with previous studies that have shown higher sorbitol levels in CSF compared to serum in healthy individuals ^25^. Sorbitol is actively transported across the blood-brain barrier ^26^ and it thus appears that the CNS and/or PNS produces and regulates levels of sorbitol independently of the periphery. Further studies are required to substantiate these hypotheses. A confirmation of higher sorbitol levels in the CSF of SORD neuropathy patients will be of interest and will potentially inform therapeutic strategies of drug delivery to the CNS.

SORD neuropathy in patients presents as moderately severe, motor dominant, axonal CMT, intermediate CMT, or dHMN ^1,27–29^. The average age of onset is the late teens ^1^. The *Sord*^−/−^ rats developed a predominantly motor neuropathy, as shown by behavioral, biochemical, electrophysiological, and especially histological studies. Behavioral studies demonstrated a decline in motor performance in the rotarod starting at ∼7 months of age; no signs of sensory deficits were demonstrated by the hotplate test. Rats developed an abnormal gait pattern detected by the MotoRater, predominantly in the hindlimbs. Electrophysiological studies revealed decreased nerve conduction velocities of peripheral nerves and normal CMAP levels. Slightly reduced NCVs have likewise been reported in patients, suggesting a loss of large myelinated nerve fibers and/or moderate de/remyelination. Histological studies of peripheral nerves in *Sord*^−/−^ rats revealed a general loss of large, myelinated fibers from the ventral root to the peripheral sections of the tibial nerve. There was mild axonal loss, and also evidence of demyelinating and remyelination, and a surprise and potentially pathognomonic finding – “ballooned” myelin sheaths. As shown by electron microscopy, these changes are localized in the compact myelin sheets, which appear split apart, potentially caused by the accumulation of fluid between the layers of myelin and the periaxonal space due to increased osmolarity mediated by high sorbitol levels. Ballooning or split fibers were previously observed in cats that developed spontaneous diabetes with neuropathy ^20^. Likewise, rats with a chronic galactose neuropathy showed similar features that were attributed to an endoneurial osmotic imbalance due to galactose intoxication ^21,22^. Galactose is also a substrate of the polyol pathway, and it is metabolized into galactitol by aldose reductase. Thus, the observed ballooned and split fibers might be a pathognomonic finding for polyol pathway alterations leading to increased osmotic pressure ^4,7^. Our findings in the *Sord*^−/−^ rats further support the hypothesis of osmolarity imbalance leading to peripheral nerve damage. Irrespective, the observed demyelination in *Sord*^−/−^ rats was unexpected and needs to be confirmed in patients.

CMT is a length-dependent neuropathy affecting distal sections of peripheral nerves. The MotoRater studies confirmed that hind limbs are predominantly affected. However, the detailed histological work up showed that the degenerative process is not exclusively length-dependent, but motoric (ventral) nerve roots show abnormal fibers compared to age matched control animals. A sensory phenotype was not observed in behavioral studies, yet histological examination shows evidence of mild pathology of sensory nerves. This is in agreement with some SORD neuropathy patients, who exhibit both motor and mild sensory symptoms. Finally, *Sord*^−/−^ rats had abnormally increased levels of serum NfL, a biomarker of axonal degeneration. This observation is valuable for preclinical studies and is potentially translatable to human studies.

Importantly, sorbitol toxicity has been implicated in the pathogenesis of diabetic neuropathy. Previous studies showed that hyperglycemia leads to sorbitol accumulation in Schwann cells, resulting in their de-differentiation, which could be rescued both *in vitro* and *in vivo* with aldose reductase inhibitors ^30^. In diabetic neuropathy, sorbitol levels are only mildly elevated, and clinical presentation occurs at an advanced age. On the other hand, SORD neuropathies and *Sord*^−/−^ rats are characterized by multi-fold increases in sorbitol levels, which are likely present throughout peripheral nerve development, including prenatal phases, which may explain the much earlier onset of clinical pathology in these settings.

In summary, the *Sord*^−/−^ rat model presented herein is closely resembling the human SORD neuropathy phenotype and has already led to new insights into disease pathophysiology. These animals represent valuable tools for further mechanistic and translational studies for diseases of high sorbitol and specifically the common hereditary SORD neuropathy.

## Supplementary material

**Supplementary Figure 1:** Genomic sequence of 18,219bp deletion at genomic coordinates (chr3:114,188,869 and chr13:114,207,088) in the *Sord*^−/−^ rat.

**Supplementary Figure 2:** Serum sorbitol measured in 12 animals (6 males and 6 females) at three different time points (baseline, first follow-up under high glucose diet (1 week after starting diet), second follow-up under diet (2 months after starting diet). One-way ANOVA, Turkey posttest for alpha correction.

**Supplementary Figure 3:** Behavioral assessment of rat motor function in different age groups with rotarod (A-D) and narrow bean (E-H) tests.

**Supplementary Figure 4: Loss of large axons in the sciatic and tibial nerves of *Sord*^−/−^ rats.** Images of semi-thin sections of the proximal sciatic nerve, tibial nerve at the knee, and tibial nerve at the ankle, from *Sord*^−/−^ and wild type (WT), as indicated. All *Sord*^−/−^ nerves show an overall loss of large fibers, and contain abnormal myelinated axons, including degenerating myelinated sheaths (arrows), thinly myelinated axon (arrowheads), and clusters of regenerated axons (circled). These changes are apparent at all levels but appear more pronounced in the distal sections of the nerve. Scale bar = 20 µm.

**Supplementary Figure 5: Myelinated sensory axons are largely preserved in the caudal nerves of *Sord*^−/−^ rats.** Images of semi-thin sections of the ventral caudal nerve from *Sord*^−/−^ (B and D) and wild type (WT; A and C) rats, as indicated. A few degenerating myelinated sheaths are present (arrows) in the *Sord*^−/−^ nerves. Scale bars = 20 µm.

**Supplementary Video:** The video shows 15 months old WT (left) and *Sord*^−/−^ (right) rats walking through the MotoRater channel (top, bottom and side views). The skeleton view illustrates the hindlimb joint points.

