## Supplemental Figures for "Sord deficient rats develop a motor-predominant peripheral neuropathy unveiling novel pathophysiological insights"

#### Supplementary Figure 1

F0 #16 (M)— DOB 8/12/2020 18219bp deletion at genomic coordinates (12743-30961)

Deletion Start
(AGTAGGGTATGGTGTGAACTTGTGTTCTGAGAAATAATTCAACGCAACTACA
GG
AGAATATTTTATAAGTGAAAAATATATAGATAAATTATTATGCAGACTAAACAAC
TTT
CTCTGTGTTATTTATTTCAGGAGAACTACCCAATCCCTGAGCTGGGCCCAAATG
G TAGGTCCTGATGTGACATTCATTGTATCTGCTGGTGTTGGCCTTGGAGGCGT
…)

Deletion End
(…CCAACAGTGCTCATTCATGTTCAGGAGCAGTACTGGTTGCTAAGCAACCAG
GAGACTTCCACCCAAAGATCCTAAATCCAGCCTAACTCATACAAGAGGGCCAC
AGGAGGGCTTGAGTTTCCCACTCACAGGATTCGCCTCCTCCCAGGCTCAC
TCCTAGGCAATTATTATCCCATCCCACTCAGAAGATGCTCCCCTTCTCGGCTG)

### Supplementary Figure 2

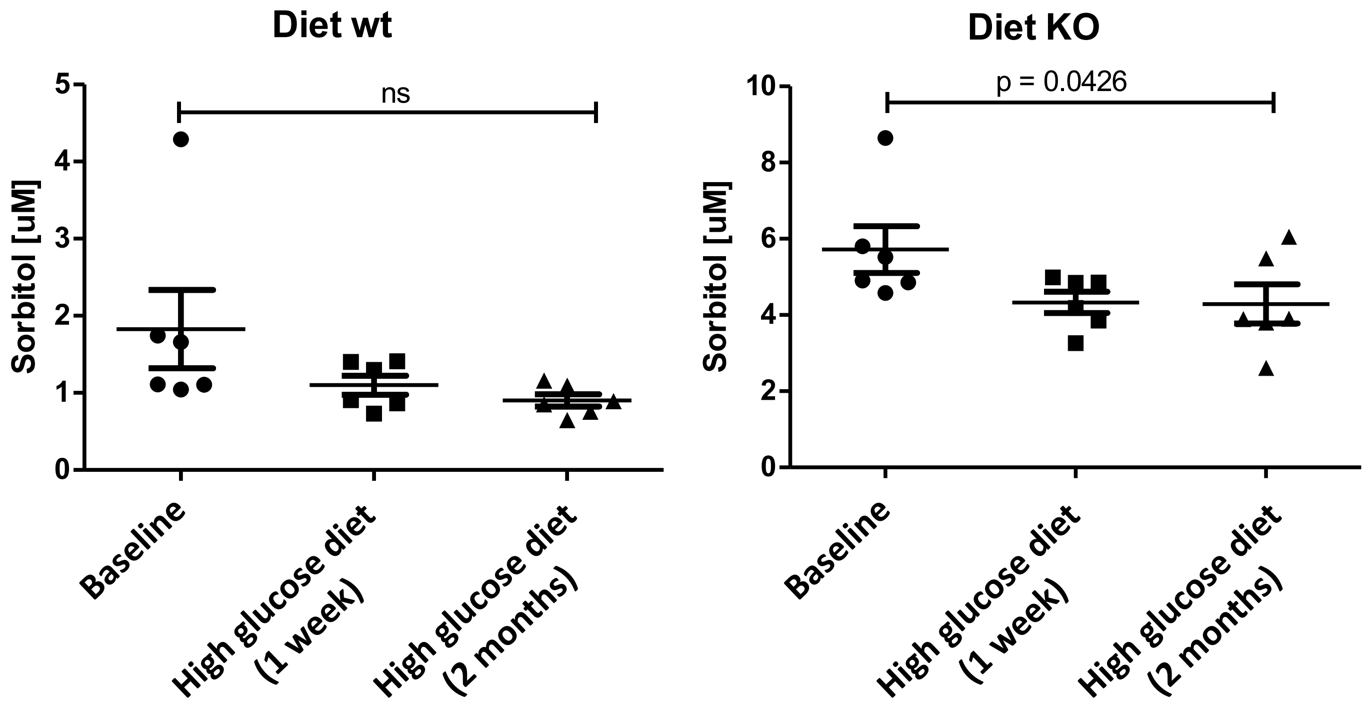

#### Supplementary Figure 3

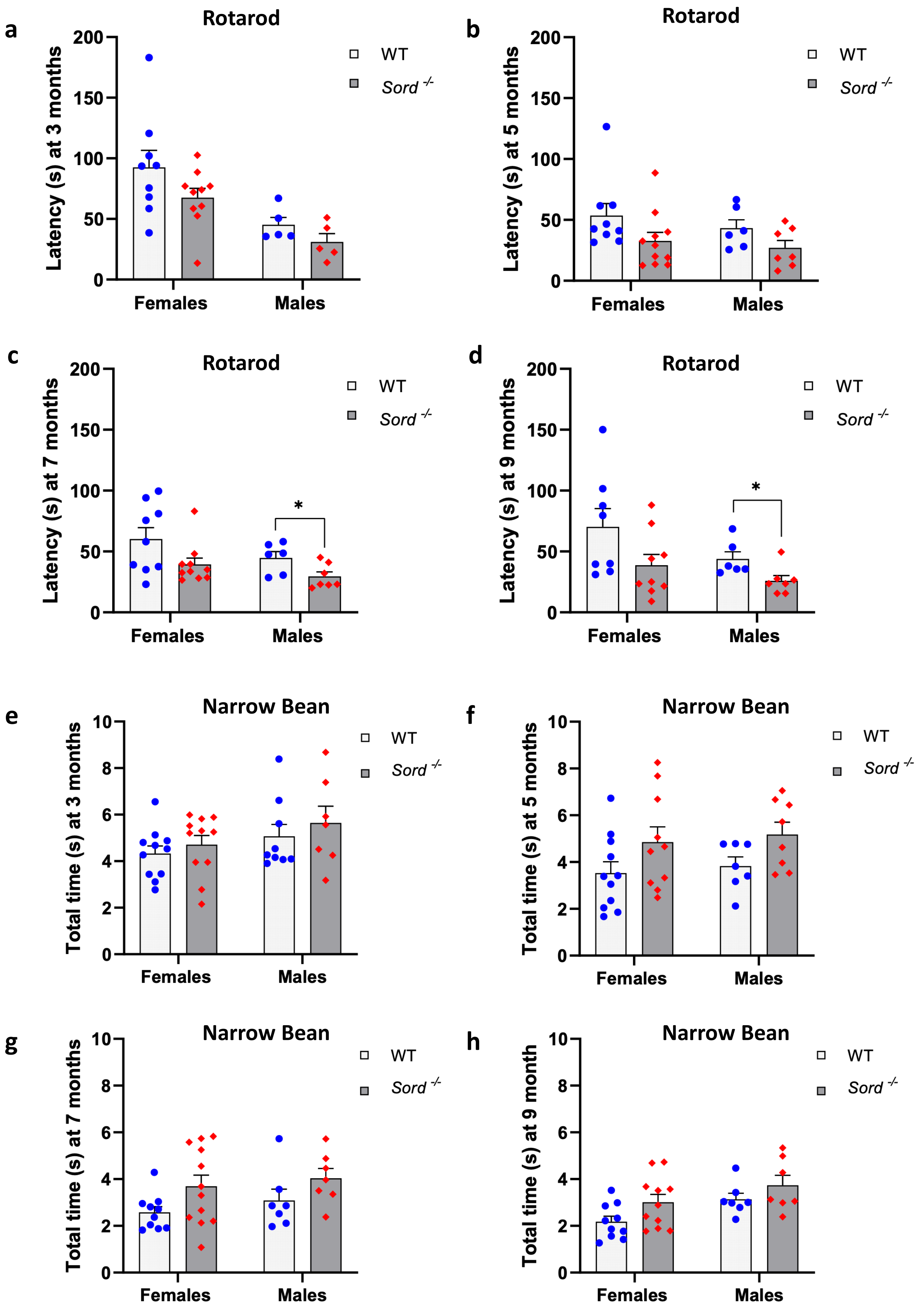

### Supplementary Figure 4

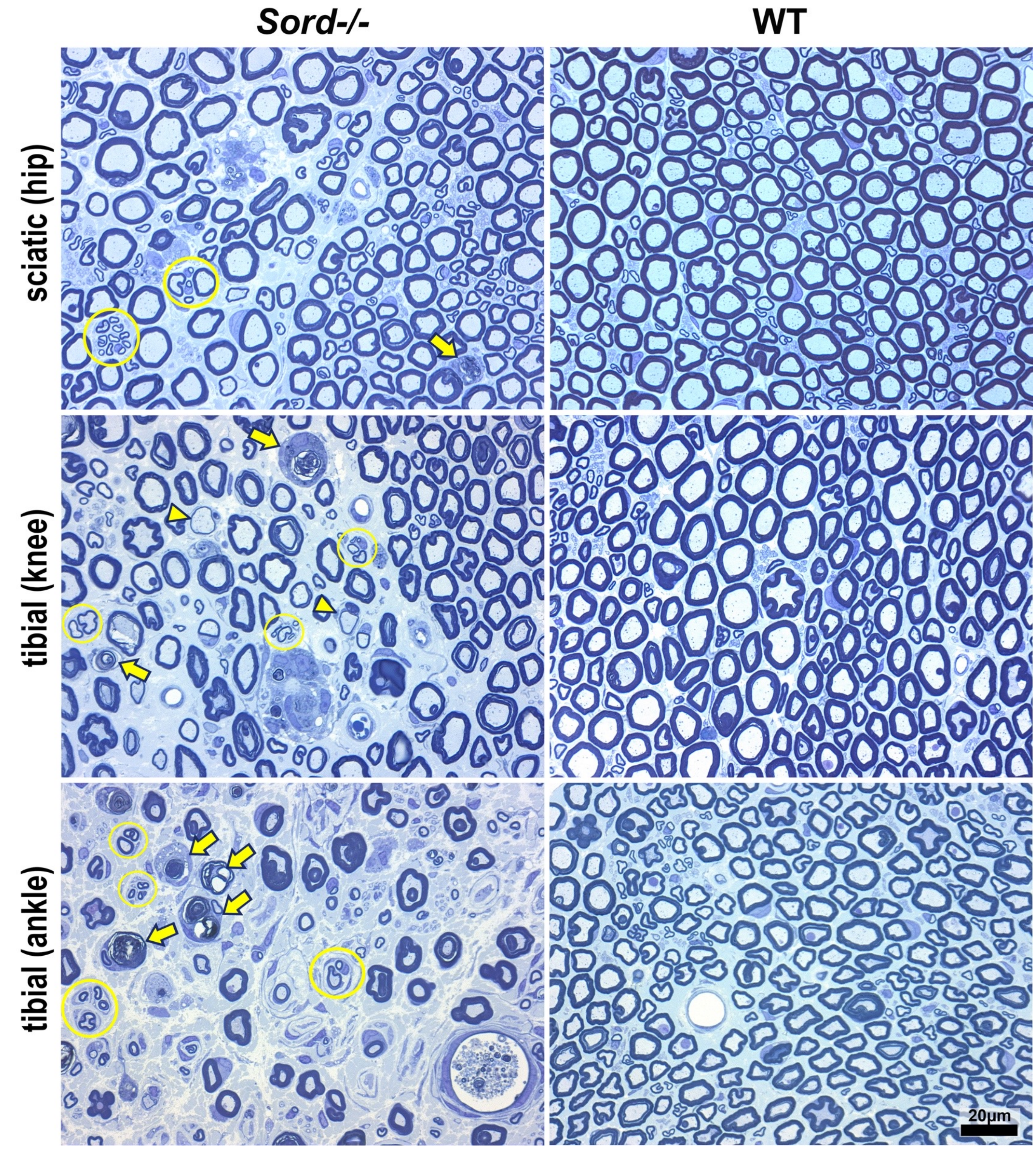

### Supplementary Figure 5

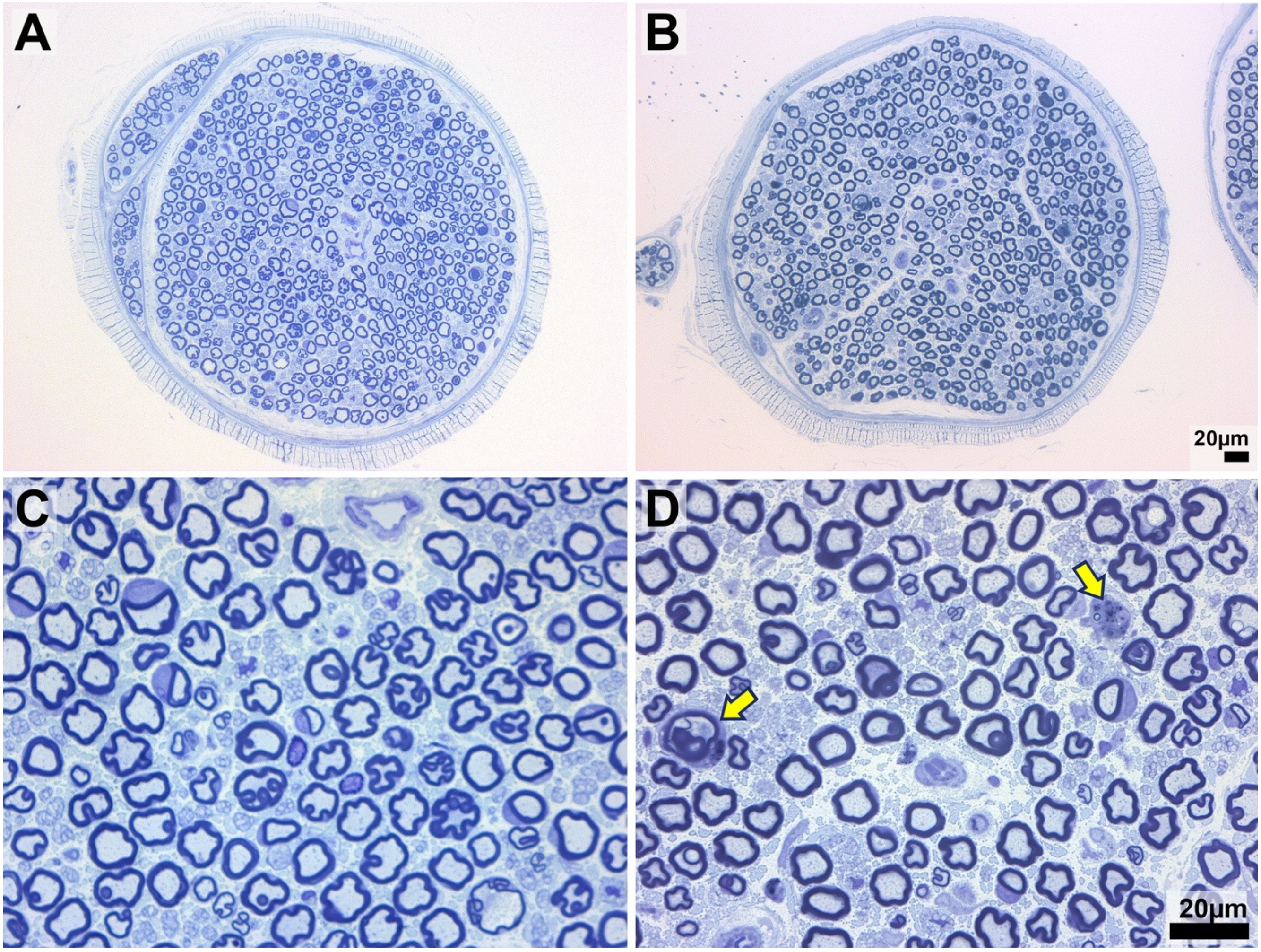
